# Excluded Volume Effect-induced Shrinkage for Resolution Enhancement of Hydrogel Bioconstructs

**DOI:** 10.64898/2026.10.25.761631

**Authors:** Dezhi Zhou, Hui Xu, Binhan Li, Bangrui Li, Shuang Yu, Liliang Ouyang

**Affiliations:** Department of Mechanical Engineering, Tsinghua University, Beijing, 100084, China; State Key Laboratory of Tribology in Advanced Equipment, Tsinghua University, Beijing, 100084, China; Beijing Key Laboratory of Intelligent Organ Biofabrication and Regenerative Repair, Tsinghua University, Beijing, 100084, China; Biomanufacturing and Engineering Living Systems Innovation International Talents Base (111 Base), Tsinghua University, Beijing, 100084, China

**Keywords:** biofabrication, hydrogel dehydration, macromolecular crowding, post-processing, resolution enhancement

## Abstract

Biofabrication of hydrogel materials is usually constrained by process-specific resolution limits. Here, we introduce excluded volume effect-induced shrinkage (EVER), a dehydration-enabled resolution enhancement strategy that achieves dimensional refinement via post-processing. In this approach, we treat biofabricated constructs with hydrophilic macromolecular crowding agent bath, with poloxamer as a representative. With EVER treatment, gelatin-based constructs can reach a dimension reduction of up to ∼49% with an estimated volumetric reduction of ∼87%. Shape fidelity was well retained after shrinkage across different constructs and could be stabilized by introducing secondary crosslinking. In extrusion bioprinted cell-laden constructs, myoblasts retained a considerable cell viability right after the shrinkage treatment and recovered significantly by day 1, with subsequent cell spreading and increasing proliferation over 7 days. Without the need of continued immersion in the shrinkage bath, EVER provides a hardware-independent route for refining hydrogel structures while retaining encapsulated cell compatibility.

## 1. Introduction

Biological tissues are organized across multiple length scales. Microscale features regulate cell alignment, migration, and collective organization, whereas macroscale architecture supports transport and mechanical integration^1^. There is a great demand for fabrication methods capable of defining fine features^2^. Specific biofabrication process carries an intrinsic resolution floor set by its physical principle, such as the nozzle diameter and filament fusion in extrusion printing^3^, the voxel size and optical scattering in light-based processing^4^, and the droplet size in inkjet printing^5^. Hardware and process optimization can contribute to resolution enhancement, but often compromises in throughput, material choice, or cytocompatibility.

Hydrogels are highly hydrated polymer networks whose swelling and shrinkage can be tuned to control their dimensions^6^. This property suggests a complementary route to finer structures: fabricating a construct at a readily accessible resolution first and then reduce its dimensions post fabrication. Previous studies have demonstrated that hydrogel architectures can be miniaturized by reducing their water content. For example, implosion fabrication uses the controlled collapse of patterned hydrogel scaffolds to produce three-dimensional nanostructures, but the required processing conditions are incompatible with living constructs^7^. Thermoresponsive pNIPAM-containing networks can shrink near physiological temperature, but it needs additional material supplements to enhance manufacturing capacity and cytocompatibility^8,9^. Electrostatic complexation can also reduce hydrogel dimensions under biocompatible conditions, but besing restricted to oppositely charged polyionic systems^10,11^. There is a great demand for versatile post-processing approach that is biocompable and can be applied across a broad range of hydrogel formulations^12^.

Macromolecular crowding provides a potential physical basis for such post-fabrication shrinkage. Concentrated polymer solutions can compact hydrated biopolymer networks through the coupled effects of dehydration^13^ and macromolecular crowding^14^, whereby solvent efflux and preferential exclusion increase the density of network-forming macromolecules and favor their intermolecular association. We reasoned that immersing a hydrated, preformed polymer network in a concentrated bath of a hydrophilic macromolecule could similarly promote water efflux until balanced by the resistance of the compacted network, resulting in a reduction in construct volume.

Here, we establish a poloxamer-based excluded volume effect-induced shrinkage (EVER) approach for enhancing the resolution of hydrogel bioconstructs, where poloxamer acts as the molecular crowding agent. Using constructs fabricated by two representative and fundamentally different routes, mold casting and extrusion-based 3D printing, we characterized the effects of poloxamer crowding bath concentration, treatment time, and hydrogel content on dimensional change. We verified that the transformation is nearly isotropic across several geometries, and distinguished reversible rehydration from dimensional stabilization by secondary photocrosslinking. To mitigate the potential effects of network compaction on encapsulated cells, we adopted a complementary-network bioink comprising a temporary gelatin network for printability and a low initial concentration of photocrosslinkable GelMA^15^. We then assessed the viability, spreading, and proliferation of encapsulated cells after shrinkage treatment. These experiments support the feasibility of applying EVER to cell-laden constructs under the tested conditions. Together, these results support a physically driven post-processing strategy in which an external crowding bath controls construct compaction. EVER does not require hardware modification or the incorporation of dedicated responsive components and is pontential to be extended to more fabrication methods.

## 2. Materials and methods

### 2.1 Materials

Gelatin methacryloyl (GelMA) was synthesized by reacting gelatin (300 g Bloom, type A, porcine skin; Sigma) with methacrylic anhydride following our previously reported protocol^16^. Gelatin, poloxamer 407 (also known as Pluronic^®^ F127) (Sigma), and lithium phenyl-2,4,6-trimethylbenzoylphosphinate (LAP; Macklin) were used as received. Poloxamer baths and all hydrogel formulations were prepared in phosphate-buffered saline (PBS) at the specified concentrations. For fixation experiments, LAP was included in the poloxamer bath at 0.5 wt%.

### 2.2 Fabrication of hydrogel constructs

Disc-shaped and patterned constructs were prepared by casting gelatin or GelMA precursor solutions into polydimethylsiloxane molds defining discs, triangles, semicircles, and blossom-shaped geometries. The filled molds were cooled at 4 °C to establish a primary physical network before EVER treatment.

### 2.3 Post-processing shrinkage

After fabrication, constructs were fully immersed in poloxamer aqueous solutions of specified concentrations at 4 °C. Dimensional changes were monitored by microscopy. The dimension ratio was defined as L/L_0_, where L is the in-plane diameter or vertical height at a given time point and L_0_ is the corresponding initial dimension. The EVER-treated constructs were either transferred directly to fresh PBS (rehydration mode) or exposed to a 405 nm UV-light (50 mW cm^−2^, 20 s) for photocrosslinking (fixation mode), followed by transferring to PBS or culture medium.

### 2.4 Shape-fidelity quantification

Binary masks were applied to the microscopy images of hydrogel constructs. For the disc-shaped constructs, the normalized roundness is calculated as R/R_0_, where R is the roundness at the indicated state and R_0_ is the initial roundness. For the complex geometries, after scale normalization, shape fidelity was characterized using the Mean boundary error, Dice coefficient, and Procrustes distance. The normalized area was calculated as A/A_0_, where A is the projected area at the indicated state and A_0_ is the initial projected area.

### 2.5 Cell-laden bioprinting, culture, and analysis

C2C12 cells were expanded in Dulbecco’s modified Eagle’s medium supplemented with 10% fetal bovine serum. Cells were suspended at 1 × 10^7^ cells mL^−1^ in a bioink containing 2.5 wt% GelMA, 5 wt% gelatin, and 0.15 wt% LAP. Cell-laden grids were fabricated by extrusion bioprinting with a 25G nozzle (SunP Biomaker2i). EVER-treated grid structures were immersed in 20 wt% poloxamer at 4 °C for 20 min and then subjected to UV exposure (50 mW cm^−2^, 20 s). Constructs were transferred to culture medium and maintained under standard culture conditions. Viability was evaluated by live/dead staining, and cell spreading was examined by confocal microscopy. Proliferation was assessed on days 1, 3, 5, and 7 using a CCK-8 assay (Yeasen) according to the manufacturer’s protocol. Optical density was measured at 450 nm using a microplate reader (Thermo Scientific). Optical density readout were normalized to the corresponding day-1 values.

### 2.6 Statistics

Statistical analyses were performed in GraphPad Prism 10.1.2. Quantitative data are presented as mean ± standard deviation (SD). Two-group comparisons used two-tailed Mann-Whitney U tests, and comparisons involving more than two groups used Welch’s analysis of variance followed by two-tailed Dunnett’s T3 multiple-comparison tests, as indicated in the figure legends. P < 0.05 was considered statistically significant; ns, not significant.

## 3. Results

### 3.1 Fundamentals of shrinkage

Poloxamer is a biocompatible, water-soluble neutral polymer long used in pharmaceutical and biomedical applications^17^ and was selected as the macromolecular crowding agent in this study. The concentration of the crowding polymer bath is expected to influence the driving force for shrinkage. We first tested whether concentrated poloxamer baths could dehydrate preformed disc-shaped hydrogels. Cast constructs were incubated in PBS or 5–40 wt% poloxamer at 4 °C for up to 12 h (**Fig. 1a, b**). PBS controls swelled progressively, whereas 5 wt% poloxamer largely maintained the initial dimensions and 10 wt% or higher poloxamer groups lead concentration-dependent shrinkage. Dimensional reduction was evident within 20 min and continued over the following 1–2 h before approaching a plateau. Quantification showed overall decreases in both in-plane and vertical dimension ratios (**Fig. 1c, d**). At 10 wt%, the in-plane ratio decreased to approximately 0.89 at 2 h and 0.84 at 12 h. At concentrations of 20 wt% or higher, the ratios at 12 h converged near 0.71–0.72 in-plane and 0.69–0.72 vertically, equivalent to approximately 30% linear and 65% volumetric reduction. The similar in-plane and vertical responses indicate near-isotropic compaction under these conditions, with a modestly greater through-thickness response at the highest concentrations, which may reflect differences in water transport and network deformation.

**Fig 1.**
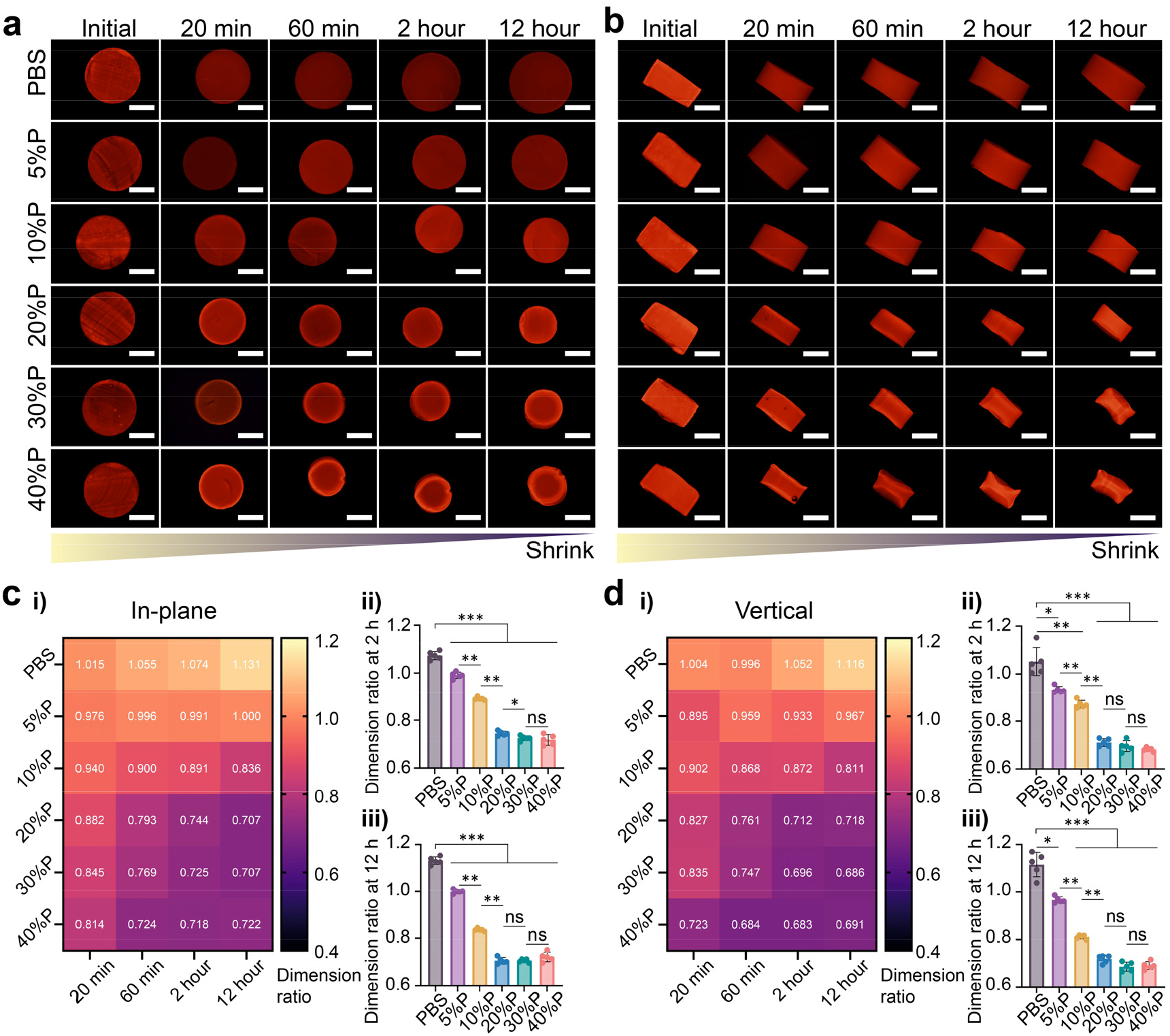
Concentration- and time-dependent shrinkage of cast Gelatin constructs. **a-b**, Representative top-view (a) and side-view (b) images of 10wt% gelatin constructs incubated at 4 °C in PBS or 5–40 wt% poloxamer (X % P) for up to 12 h. **c-d**, Heat maps and quantification of the in-plane (c) and vertical (d) dimension ratios. Data are presented as mean ± SD (n = 5). Welch’s ANOVA followed by two-tailed Dunnett’s T3 multiple-comparison tests was used for comparisons with the PBS control; two-tailed Mann-Whitney U tests were used for two-group comparisons without PBS control. ns, not significant; *P < 0.05, **P < 0.01, ***P < 0.001. Scale bars: 2 mm.

When shrunk constructs were transferred directly to PBS, they progressively rehydrated toward their initial in-plane and vertical dimensions (**Supplementary Fig. 1**). This reversibility supports a predominantly physical dehydration-rehydration process. The disc overall well retain its integrity during shrinkage, although there will be slight deformation for the constructs when the crowding agent concentration goes too high (e.g., 40 wt%) (**Fig. 1a,b**). We therefore selected 20 wt% poloxamer as the standard condition for subsequent experiments.

### 3.2 Polymer-content dependence and fixation of shrunk constructs

Unfixed constructs reabsorbed water and swelled toward their initial dimensions in PBS (**Supplementary Fig. 1**). To stabilize the shrunk state, we introduced a secondary photocrosslinking network. We select GelMA for structure fabrication as it supports both physical gelation upon cooling and covalent crosslinking upon light exposure. Because the matrix polymer content influences both water retention and network resistance to compaction, we next examined its effect on the attainable dimensional reduction. GelMA constructs of 3–50 wt% were incubated in 20 wt% poloxamer bath (**Fig. 2a**). The dimension ratio increased with GelMA content (**Fig. 2b**). At 12 h, 3 wt% GelMA reached ∼0.51, corresponding to a 49% linear reduction and an estimated 87% volumetric reduction. By contrast, 50 wt% GelMA remained close to its initial size (dimension ratio ∼1.02), and formulations containing 5–30 wt% GelMA showed intermediate responses. Lower concentration of GelMA present more removable water and lower resistance to compaction in the crowding agent bath, whereas polymer-rich networks provide less water and higher mechanical resistance.

**Fig 2.**
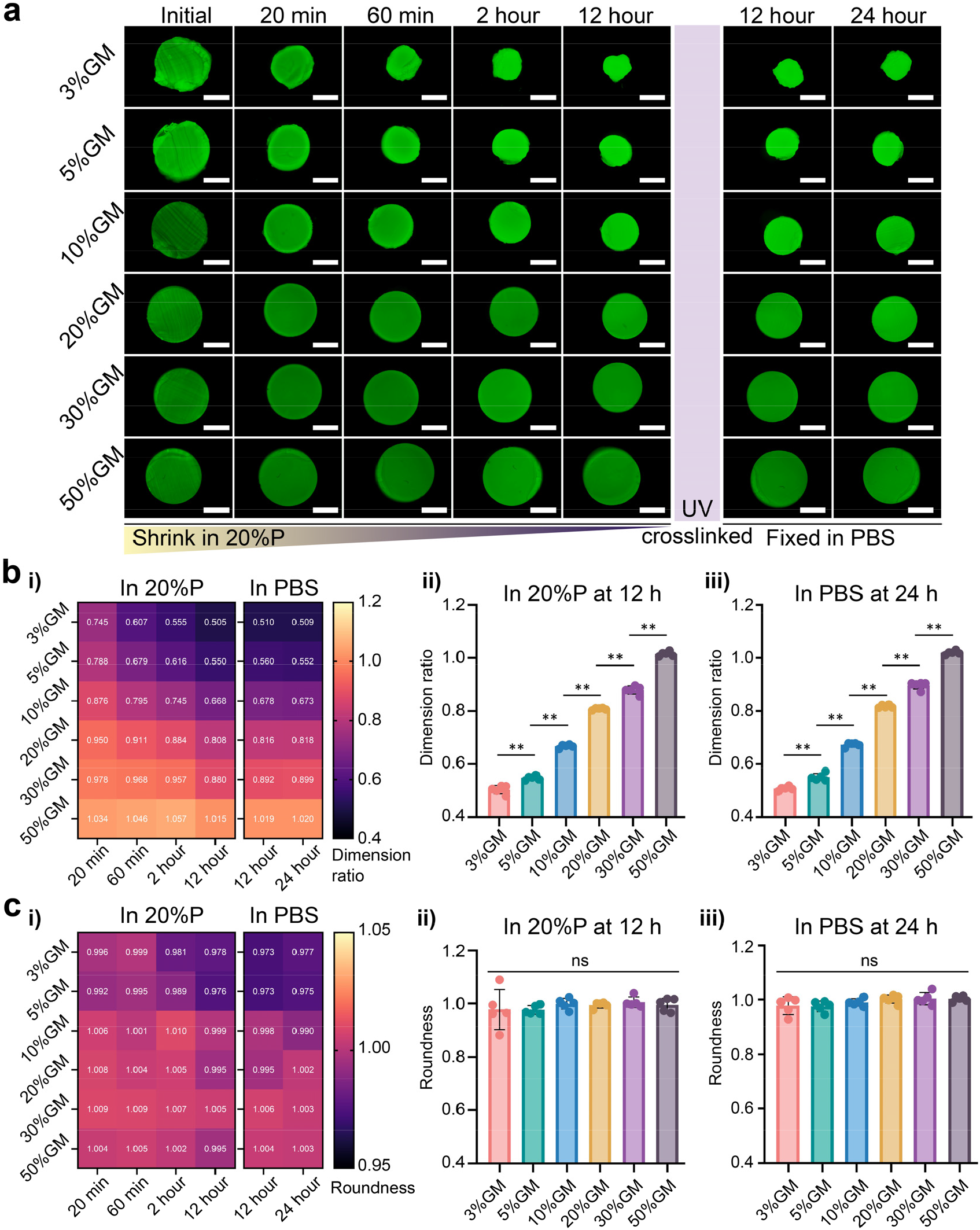
Polymer-content dependence and fixation of compacted GelMA constructs. **a**, Representative images of constructs containing 3–50 wt% GelMA during incubation in 20 wt% poloxamer (20%P) at 4 °C, after photocrosslinking after dehydration, and after transfer to PBS. **b-c**, Heat maps and quantification of the dimension ratio (b) and normalized roundness (c) during dehydration and after fixation in PBS. Data are presented as mean ± SD (n = 5). Two-tailed Mann-Whitney U tests were used for two-group comparisons, and Welch’s ANOVA followed by two-tailed Dunnett’s T3 tests was used for multiple comparisons. ns, not significant; *P < 0.05, **P < 0.01, ***P < 0.001. Scale bars: 2 mm.

Because unfixed constructs rehydrated in PBS, we used secondary photocrosslinking to retain the compacted dimensions. Constructs were crosslinked while still immersed in the poloxamer bath and were then transferred to PBS. The dimensions remained stable for 24 h across the tested formulations; for example, 3 wt% GelMA exhibited a shrinkage dimension ratio of 0.505 after 12 h shrining in poloxamer and maintained the ratios of 0.510 and 0.509 after 12 and 24 h rehydration in PBS, respectively (**Fig. 2b**). EVER can therefore be used in a reversible rehydration mode or combined with secondary crosslinking to produce a fixed state. Normalized roundness remained near 1.00 during dehydration and after fixation in PBS (**Fig. 2c**), indicating that circular constructs largely retained their circular shape.

### 3.3 Shape fidelity across complex geometries

To determine whether EVER preserved noncircular outlines, we analyzed triangle, semicircle, and blossom-shaped GelMA constructs treated with 20 wt% poloxamer. The overall outlines were retained after dehydration and after photocrosslinking followed by 24 h in PBS (**Fig. 3a**). After scale normalization, the initial, dehydrated, and fixed-in-PBS contours of the blossom geometry showed close superposition (**Fig. 3b**). Mean boundary error, Dice coefficient, and Procrustes distance did not differ significantly between the dehydrated and fixed-in-PBS states for any of the three geometries (**Fig. 3c**). The normalized projected areas clustered near A/A0 = 0.33 in both states and did not differ significantly among the tested shapes (**Fig. 3d**). Thus, within this set of geometries, EVER produced a reproducible dimensional reduction while preserving shape fidelity, and secondary crosslinking maintained the compacted geometry in PBS.

**Fig 3.**
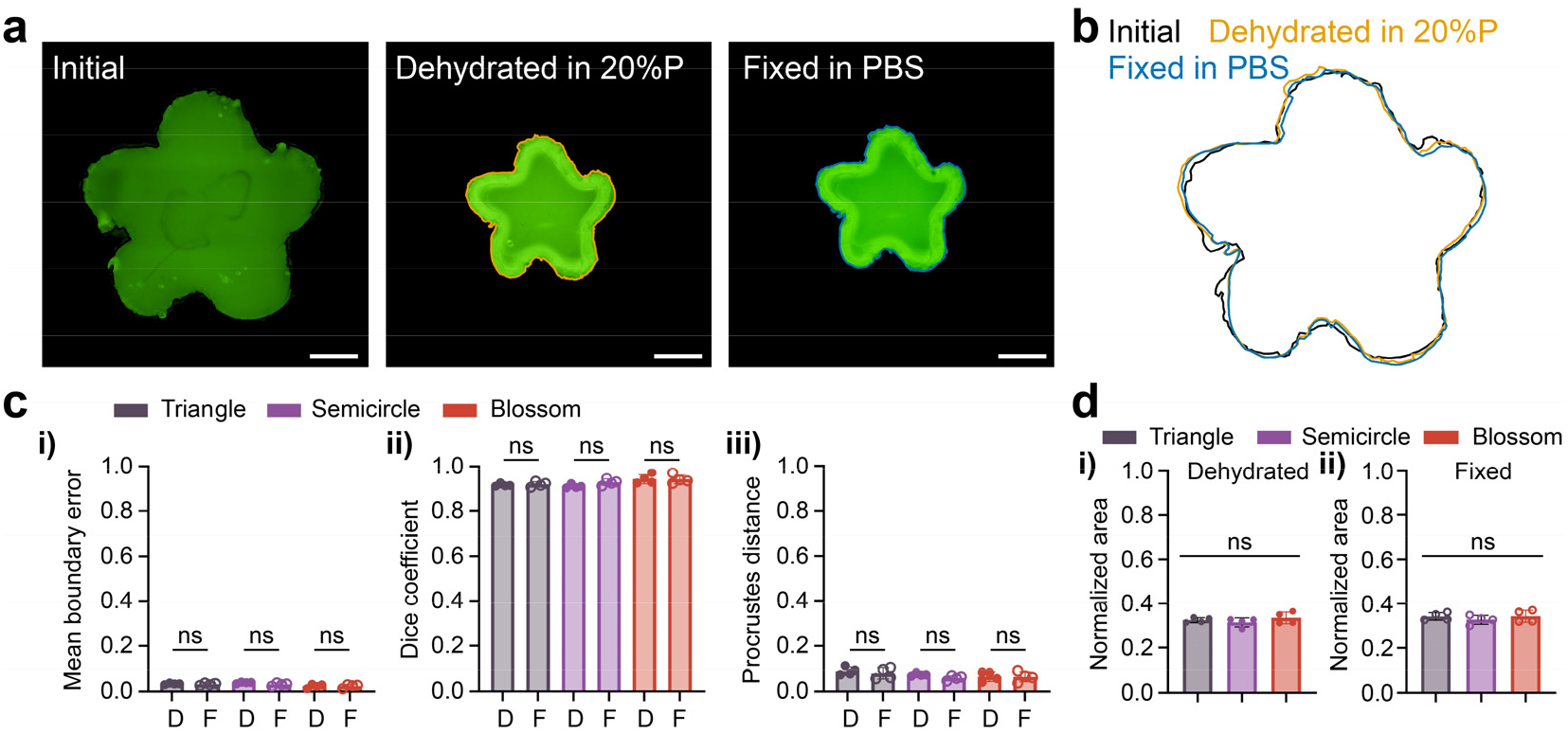
Shape fidelity of complex geometries during dehydration and after fixation in PBS. **a**, Representative images of a blossom-shaped GelMA construct in the initial state, after 24 h in 20 wt% poloxamer at 4 °C (dehydrated), and after photocrosslinking followed by 24 h in PBS (fixed in PBS). **b**, Overlay of the scale-normalized contours of the three states. **c**, Mean boundary error, Dice coefficient, and Procrustes distance for triangle, semicircle, and blossom geometries in the dehydrated (D) and fixed-in-PBS (F) states. **d**, Normalized projected area (A/A0) for the three geometries in the dehydrated and fixed-in-PBS states. Data are presented as mean ± SD (n = 5). Two-tailed Mann-Whitney U tests were used for two-group comparisons, and Welch’s ANOVA followed by two-tailed Dunnett’s T3 tests was used for multiple comparisons. ns, not significant; *P < 0.05, **P < 0.01, ***P < 0.001. Scale bars: 2 mm.

### 3.4 Cytocompability

Cell printing enables the direct deposition of cells into 3D constructs. However, the incorporation of living cells introduces additional constraints on fabrication resolution: smaller nozzles will compromise cell viability in extrusion- and jet-based processes, whereas high cell densities can increase light scattering in optical printing. A cell-compatible dehydration step may therefore offer a complementary route to refining construct dimensions after fabrication. Here, we evaluated whether EVER treatment was compatible with living cells using extrusion-printed myobalst-laden grids as a proof of concept. Grids containing 2.5 wt% GelMA and 5 wt% gelatin were printed through a 25G nozzle, immersed in 20 wt% poloxamer at 4 °C for 20 min, and then UV-crosslinked for culture (**Fig. 4a**). Immediately after processing, viability in the EVER group was approximately 68%, which was a bit lower than non-dehydrated controls (∼80%) (**Fig. 4b**). By day 1, viability increased to approximately 79% and was no longer significantly different from the control. Electrostatic approaches use oppositely charged polymers to induce shrinkage^10,11^, while our approach uses transient exposure to a poloxamer bath. Constructs are transferred to culture medium after fixation without requiring continued immersion in the bath, with less persistent influences on cells.

**Fig 4.**
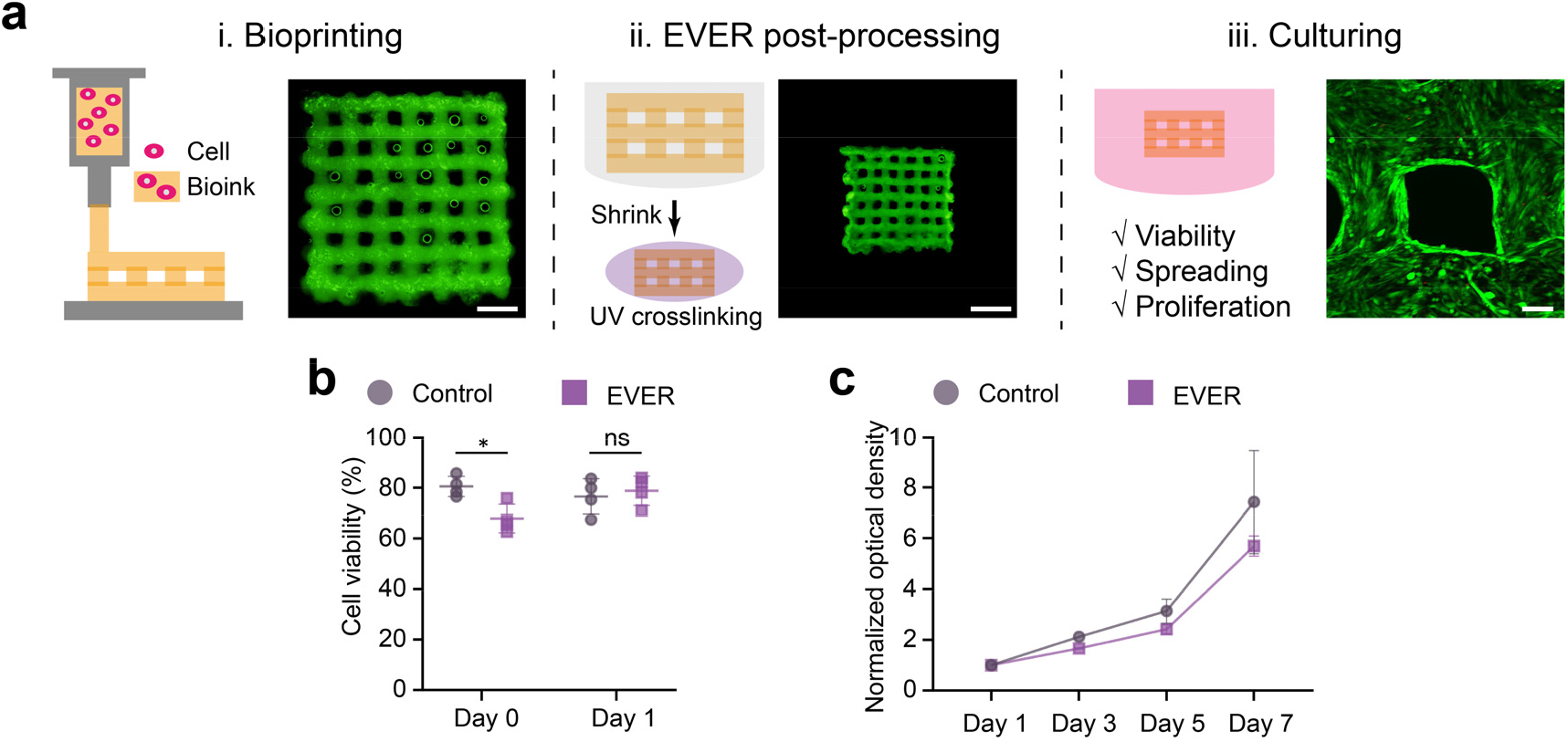
Cell response to EVER post-processing of extrusion-printed constructs. **a**, Workflow comprising extrusion printing of myoblast-laden grids, EVER treatment in 20 wt% poloxamer at 4 °C for 20 min, photocrosslinking, and subsequent culture. Representative images show the grid structure before and after treatment and cell spreading after 7-day culture. **b**, Viability in non-dehydrated control and EVER-treated constructs at days 0 and 1. **c**, CCK-8 assay of proliferation over 7 days, expressed as normalized optical density relative to day 1. Data are presented as mean ± SD (n ≥ 3). Two-tailed Mann-Whitney U tests were used for two-group comparisons. ns, not significant; *P < 0.05, **P < 0.01, ***P < 0.001. Scale bars: 2 mm (i and ii) and 200 µm (iii).

Cells in the compacted constructs subsequently spread within the matrix, as observed by confocal imaging (**Fig. 4a**). CCK-8 based proliferation-fold increased over 7 days in both groups, with the EVER-treated constructs reaching approximately 5–6 times of their day-1 data (**Fig. 4c**). This result supports continued proliferation-associated metabolic activity after EVER processing. Volumetric shrinkage can increase polymer concentration and alter matrix mechanics^9,10^, potentially affecting encapsulated cells^12^. To balance printability with a cell-supportive matrix after processing, we adopted a complementary-network strategy^15^ in which gelatin provides temporary physical support, allowing the use of a low initial GelMA concentration. During culture, the gelatin component is expected to dissolve and diffuse out, leaving the photocrosslinked GelMA network to support the construct. This complementary-network design may help separate the material requirements for printing from those for cell culture after compaction. Provided that cells are retained during processing, volumetric compaction is expected to increase their spatial density immediately, while subsequent proliferation may further increase cell density if construct volume remains stable. EVER approach may therefore offer a route to coupling geometric miniaturization with cellular densification in engineered tissues.

### 3.5 Discussion and conclusions

This study establishes a post-fabrication route for reducing the dimensions of fabricated hydrogel constructs. Using manufacturing techniques such as mold casting or extrusion printing defines the initial architecture, whereas a subsequent poloxamer bath controls compaction. The successful application of EVER to both cast and extrusion-printed constructs supports its compatibility with the two fabrication routes tested here, with potential extension to other printing methods and material classes. This sequential design separates architecture formation, dimensional refinement, and stabilization, allowing each step to be controlled through a distinct processing variable.

Our approach differs from existing shrinkage-based approaches in the mechanism and applicable materials. Implosion fabrication achieves greater miniaturization degree but relies on harsh dehydration of synthetic scaffolds^7^. pNIPAM-based systems offer thermally triggered shrinkage but require thermoresponsive components and exhibit formulation-dependent cell interactions^8,9^. Electrostatic complexation can be cytocompatible and highly effective, but requires oppositely charged networks and polyions, with cell tolerance depending strongly on polycation concentration and cell type^10,11^. In EVER an external, water-soluble poloxamer bath compacts a preformed physical network, after which crosslinking (e.g. photocrosslinking) stabilizes the compacted structure. This sequence avoids the need to incorporate dedicated thermoresponsive components or oppositely charged polymer pairs. The complementary-network formulation further offers a way to balance manufacturing capacity with the matrix requirements of cells after compaction.

The polymer concentration and type dependence, near-isotropic dimensional change, and reversibility are consistent with a crowding-associated reduction in water activity outside the hydrogel, followed by water efflux until the dehydration driving force is balanced by the resistance of the compacted network. This interpretation is consistent with the rapid initial response and subsequent plateau.

The cell experiments define both the promise and the present biological boundary of EVER. A 20-min treatment at 4 °C result in a considerable decrease in myoblast viability, followed by recovery by day 1 and cell spreading, and sustained proliferation over 7 days. These findings support the feasibility of EVER for cell-laden constructs. Future studies should resolve cellular responses to dehydration and altered matrix mechanics, directly quantify cell-density changes, and determine whether phenotype and function are maintained in primary or more sensitive cell types.

There remain some technical limitations as well. Dehydration is diffusion-dependent and may result in spatial heterogeneity in larger constructs, which is the general issue for all dehydration-based approaches. Although the a majority of cells are alive, it is to be investigated whether the cells would shrink or dehydrate and how would they be influenced in terms of biological functions. Together, by shifting part of the resolution challenge from the fabrication apparatus to a controllable post-processing step, EVER offers a foundation for future studies across additional hydrogel formulations and biofabrication platforms.

## 5. Supplementary materials

**Supplementary Fig. 1.**
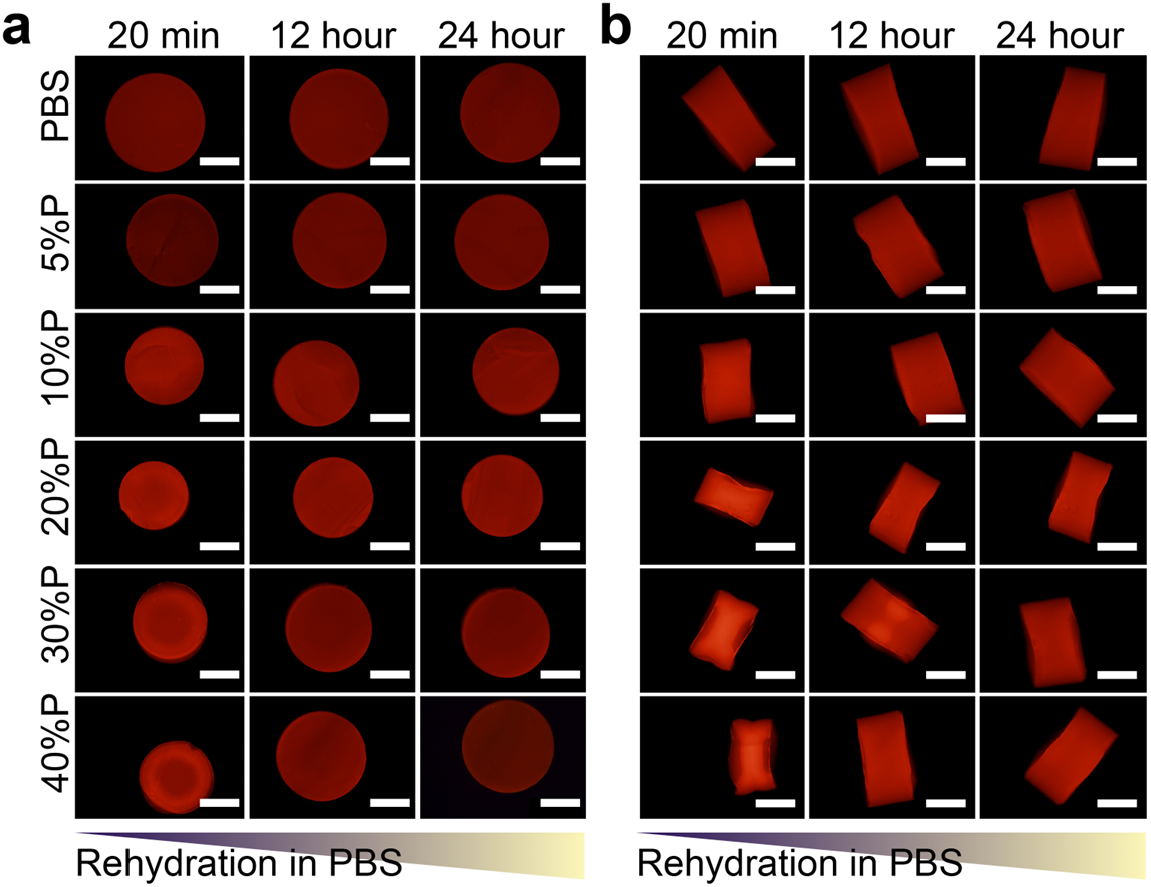
Rehydration of shrunk constructs in PBS. **a-b**, Representative top-view (a) and side-view (b) images during rehydration in PBS after constructs had been dehydrated using the indicated poloxamer concentrations for 12 hours. Images were acquired at 20 min, 12 h, and 24 h after transfer to PBS. Scale bars: 2 mm.

## Acknowledgments

The authors acknowledge B. Dou (Tsinghua University) for preparing experimental materials and Y. Lin (Beijing University of Chemical Technology) for helpful discussions. Liliang Ouyang acknowledges support from the National Natural Science Foundation of China (Nos. 52475305, U25D9019).

## Author Contributions

D.Z. and L.O. conceived and designed the study. D.Z. developed the post-processing protocol. D.Z., H.X., B.H.L., B.R.L., and S.Y. performed experiments. D.Z. analyzed the data. D.Z. and L.O. drafted the manuscript. All authors reviewed the manuscript. L.O. obtained funding and supervised the study.

